# Modular Noncovalent Functionalization of Electrospun Piezoelectric Scaffolds with Bioactive Nanocarriers

**DOI:** 10.1101/2025.10.25.684210

**Authors:** Sarah Payne Bortel, Sumayia Saif Jaima Chowdhury, Jeremy Cheng, Daniella Uvaldo, Mackenzie Wright, Treena Livingston Arinzeh, Santiago Correa

## Abstract

Electrospun scaffolds offer a promising platform for immune-instructive materials, but stable and modular functionalization with bioactive signals remains a technical challenge. Here, we develop a surface coating strategy for electrospun scaffolds that consist of poly(vinylidene fluoride-trifluoroethylene) (PVDF-TrFE), a piezoelectric polymer, using electrostatic adsorption of charged nanoparticles. We show that under certain conditions, these piezoelectric scaffolds are suitable substrates for electrostatic self-assembly, and that the density of nanoparticle coatings can be tuned by adjusting buffer pH, ionic strength, and nanoparticle concentration. This approach enables robust and uniform coating of both polymeric nanoparticles and soft nanocarriers such as liposomes, without requiring covalent surface modification. Liposome-coated scaffolds are cytocompatible with adherent epithelial and suspension immune cells and support lipid exchange at the cell-material interface. Using a supramolecular tethering strategy, we use liposome coatings to present interleukin-15 (IL-15) from the scaffold surface and demonstrate localized, sustained cytokine signaling. Together, these findings establish a modular approach for post-fabrication, noncovalent scaffold functionalization with bioactive nanocarriers, offering new opportunities for tissue and immune engineering.

## Introduction

Instructive biomaterials that modulate immune cell function are needed across regenerative medicine, cancer immunotherapy, and ex vivo cell engineering. Local delivery from a material interface can reduce systemic immune-related adverse effects,^1–6^ yet presenting potent immune signals such as cytokines in a sustained, spatially confined manner remains difficult. Cytokines are rapidly degraded in vivo,^7–9^ and often act most effectively when recognized in a membrane-proximal configuration,^9,10^ which complicates controlled use in biomaterial systems. As a result, there is a need for tunable biomaterials to locally present cytokines at the cell interface.

Prior strategies to localize bioactive cues within biomaterials include bulk encapsulation,^11^ covalent tethering,^12^ and immobilization onto nanoparticles,^13^ each with tradeoffs in spatial precision, loading efficiency, and functional potency. In soluble form, cytokines typically require repeated dosing or chemical modification to extend half-life, with associated losses in activity.^14,15^ Liposomes offer a promising alternative because they can carry fragile proteins,^16,17^ limit systemic exposure,^18^ and mimic aspects of membrane-bound ligand display.^19,20^ However, methods to integrate liposomes uniformly and stably onto scaffolds, while preserving cell access to surface-presented cargo, remain limited.^21–25^

Electrospun scaffolds are widely used in immunoengineering and tissue regeneration due to their high surface area,^26^ tunable architecture,^27–29^ and mechanical responsiveness.^30–33^ These features make electrospun scaffolds attractive for presenting bioactive signals at the cell interface, yet reliably achieving such presentation remains challenging. Addressing this gap requires a noncovalent surface functionalization approach that preserves scaffold architecture and maintains nanocarrier bioactivity and cell accessibility. We hypothesized that electrostatic self-assembly principles established in layer-by-layer systems could be adapted to electrospun fibers to tether charged nanoparticles without covalent chemistry, with coating density tuned through surface charge and solution conditions.^34,35^ Piezoelectric polymers like poly(vinylidene fluoride-trifluoroethylene) (PVDF-TrFE) present a unique opportunity in this context: mechanical deformation can induce surface charge on the fibers, creating favorable conditions for adsorption of oppositely charged species.^36–39^

Here, we present a surface-functionalized scaffold system based on electrospun PVDF-TrFE fibers, designed for tunable electrostatic adsorption of charged nanoparticles and liposomes. We show that mechanically generated surface charge during loading, together with optimized buffer conditions for pH and ionic strength, enables robust adsorption of both polymeric nanoparticles and soft lipid nanocarriers without covalent modification. The coated scaffolds remain cytocompatible and support three-dimensional cell culture of HEK and Jurkat T cells. Building on a supramolecular tethering strategy previously applied in hydrogel systems,^40^ we noncovalently display interleukin-15 on scaffold-bound liposomes and demonstrate receptor signaling in a reporter cell line. This work establishes a modular, bioactive interface for localized cytokine presentation with potential applications in *in situ* immune modulation and *ex vivo* T cell expansion.

## Results and Discussion

### 2.1 Tunable electrostatic adsorption of polymeric nanoparticles onto piezoelectric scaffolds

Electrostatic self-assembly offers a versatile strategy for surface functionalization, as exemplified by layer-by-layer (LbL) techniques that use alternating charged species to build well-defined interfaces.^41^ While these approaches have been widely applied to planar or chemically modified substrates,^42–44^ their applications to three-dimensional, electrospun fibrous scaffolds are less explored. We hypothesized that the intrinsic surface charge of PVDF-TrFE,^45^ which is a piezoelectric polymer, could support electrostatic nanoparticle adsorption without requiring covalent modification. To test this, we investigated how nanoparticle composition and coating method influence electrostatic adsorption onto these piezoelectric substrates.

To explore how nanoparticle surface properties influence scaffold adsorption, we compared two nanoparticle types: commercial carboxylate modified latex (CML) particles, which are stiff and smooth, and LbL-coated CML nanoparticles, which are expected to be softer and have a rougher surface due to their loopy, entangled polymer layers.^46^ We generated cationic LbL nanoparticles by sequentially coating CML particles with three polyelectrolyte layers: cationic poly-allylamine hydrochloride (PAH), anionic poly(acrylic acid) (PAA), and a final layer of PAH. This multilayered coating increased the particle diameter from 117.4 ± 0.1 nm to 143.6 ± 0.5 nm, as measured by dynamic light scattering. Both particle types displayed narrow size distributions (PDI = 0.03 ± 0.01 or 0.05 ± 0.01, respectively) (Supplementary Figure 1), and their high-magnitude zeta potentials (-52.00 ± 0.91 mV or +48.59 ± 2.02 mV, respectively) confirmed strong surface charge, making them suitable for electrostatic adsorption (Figure 1A).

**Figure 1.**
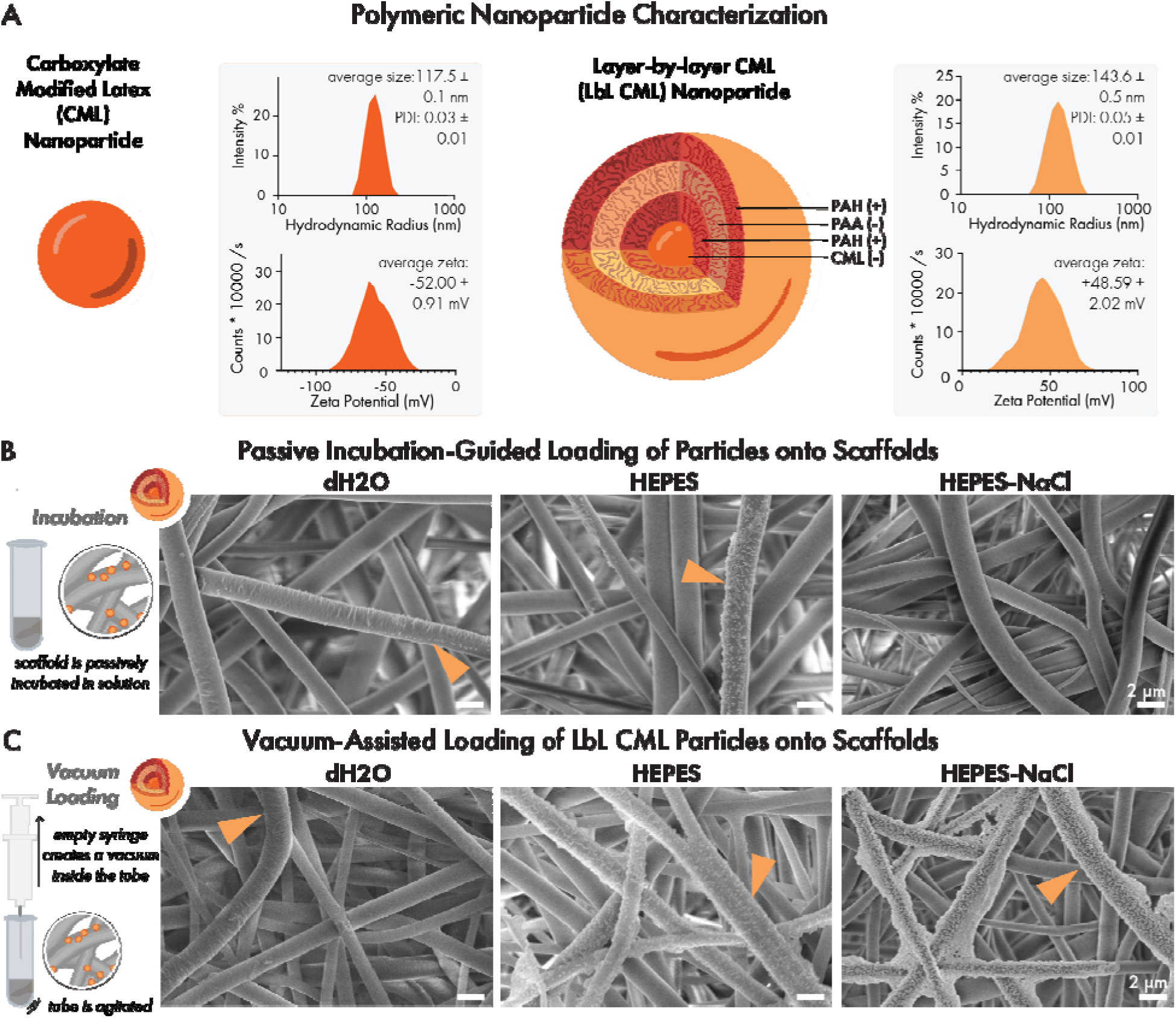
Systematic comparison of loading methodology on charged particles reveals optimal loading technique: (A) Carboxylate modified latex (CML) particle (left) and layer-by-layer CML particles (LbL CML, right). Hydrodynamic radius and zeta potential are characterized with dynamic light scattering; values are reported as mean ± SEM with n=3 replicates. (B) Scanning electron microscopy images of particle coatings with incubation-guided loading in deionized water (left), HEPES buffer (middle), or HEPES buffer + NaCl with LbL CML NPs at 2 mg/mL. (C) Scanning electron microscopy images of particle coatings with vacuum-assisted loading in deionized water (left), HEPES buffer (middle), or HEPES buffer + NaCl with LbL CML NPs at 2 mg/mL. Arrows indicate discrete particles and/or particle plaques.

To generate a suitable substrate for electrostatic adsorption, we synthesized PVDF-TrFE scaffolds and confirmed a fiber diameter of 1.80 ± 0.51 µm by scanning electron microscopy (SEM) (Supplementary Figure 2), consistent with previous reports of electrospun PVDF-TrFE scaffolds.^47,48^ To facilitate electrostatic adsorption of nanoparticles, the substrate must carry a strong surface charge of opposite polarity to the adsorbing species.^35^ Piezoelectric materials like PVDF-TrFE offer a unique advantage in this context: they can generate surface charge intrinsically through the alignment of molecular dipoles.^45,49,50^ This property can be harnessed in two ways. First, stable charge domains can be established through corona poling, in which the scaffold is exposed to a high-voltage electric field to induce long-range dipole alignment and persistent polarization.^47,51,52^ Second, mechanical deformation can induce local dipole reorientation and transient electric fields, activating surface charge in a dynamic and spatially heterogeneous manner.^53,54,49^

To optimize scaffold coating, we evaluated how these charge-generating mechanisms influence nanoparticle adsorption by comparing two nanoparticle loading strategies using corona-poled scaffolds: (i) passive incubation and (ii) vacuum-assisted deposition. For passive incubation, scaffolds were submerged in nanoparticle solution without agitation. In contrast, the vacuum-assisted method was designed to introduce mechanical stress during coating by placing scaffolds in a sealed tube filled with nanoparticle solution and applying negative pressure via syringe during repeated agitated to induce scaffold flexion. SEM revealed that passive incubation-based loading resulted in minimal nanoparticle adsorption onto the scaffold (Figure 1B), whereas vacuum-assisted loading produced significantly denser and more uniform coatings (Figure 1C).

Although we observed that mechanical activation enhanced nanoparticle adsorption, the overall coating density remained low across tested conditions. We hypothesized that, beyond surface charge effects, adsorption efficiency may also depend on solution conditions known to modulate electrostatic interactions – a well-established principle in layer-by-layer electrostatic assembly.^34,55,56^ To test this, we investigated how solution pH and ionic strength influence nanoparticle binding, with the goal of optimizing coating conditions to achieve a thicker and denser coating.

We evaluated scaffold coating in three buffer systems designed to independently modulate pH and ionic strength. Buffer pH was selected to stabilize the ionization states of charged functional groups on the nanoparticle surface. CML nanoparticles bear – COOH groups (pKa ∼4.5–5.0) that are fully deprotonated and negatively charged above pH 6, while PAH, used as the terminal layer on LbL nanoparticles, contains primary amines (pKa ∼8.5–9.3) that remain protonated and positively charged below this range. HEPES buffer (500 mM, pH 7.5, ionic strength=0.5M, Debye electrostatic screening length λ_D_=0.430 nm) was chosen to ensure strong ionization of both groups and minimize pH drift during loading. In parallel, we tuned ionic strength to regulate interparticle and particle–scaffold electrostatic interactions: low ionic strength minimizes charge screening but can increase repulsion between similarly charged species, while moderate ionic strength can reduce repulsion and promote closer packing.

To create these conditions, we used deionized water as a low-ionic strength, unbuffered condition. HEPES supplementation offered pH control with moderate ionic strength.^35^ Lastly, 500 mM HEPES supplemented with 400 mM NaCl introduced a high ionic strength (ionic strength=0.9M, Debye electrostatic screening length λ_D_=0.320 nm) to enhance adsorption. Each condition was tested at a high and low nanoparticle concentration (2 mg/mL and 0.5 mg/mL), alongside buffer-only controls. SEM revealed that when employing vacuum-assisted loading, scaffolds treated in HEPES-NaCl exhibited the most uniform and dense coverage, had moderate adsorption in HEPES alone, and were sparsely and inconsistently coated by particles in deionized water (Figure 1C). These findings indicate that optimal adsorption required simultaneous tuning of both pH and ionic strength to support charge stability and minimize repulsive interactions. However, in the passive incubation condition, modest nanoparticle coverage was observed when loading in HEPES buffer alone, but little to no coverage occurred in HEPES supplemented with NaCl (Figure 1B). We speculate that in the absence of mechanical agitation, the scaffold fibers possess low surface charge, and that the addition of salt leads to near-complete electrostatic charge screening that prevents interactions between nanoparticles and the scaffold.

We next compared the effect of particle charge on adsorption. Notably, particle adsorption was observed for both cationic and anionic nanoparticles (Figure 2 A, B), suggesting that the scaffold surface does not carry a single dominant charge. Instead, the fibrous piezoelectric architecture likely generates heterogeneous surface potentials during mechanical deformation, with coexisting microdomains of positive and negative charge. This is consistent with prior studies showing that strain gradients in piezoelectric polymers, particularly in thin and flexible materials, can produce spatially patterned electric fields.^57^ Several groups have leveraged this behavior to guide polymer adsorption or create self-patterned materials using electrostatic self-assembly principles.^58–60^

**Figure 2.**
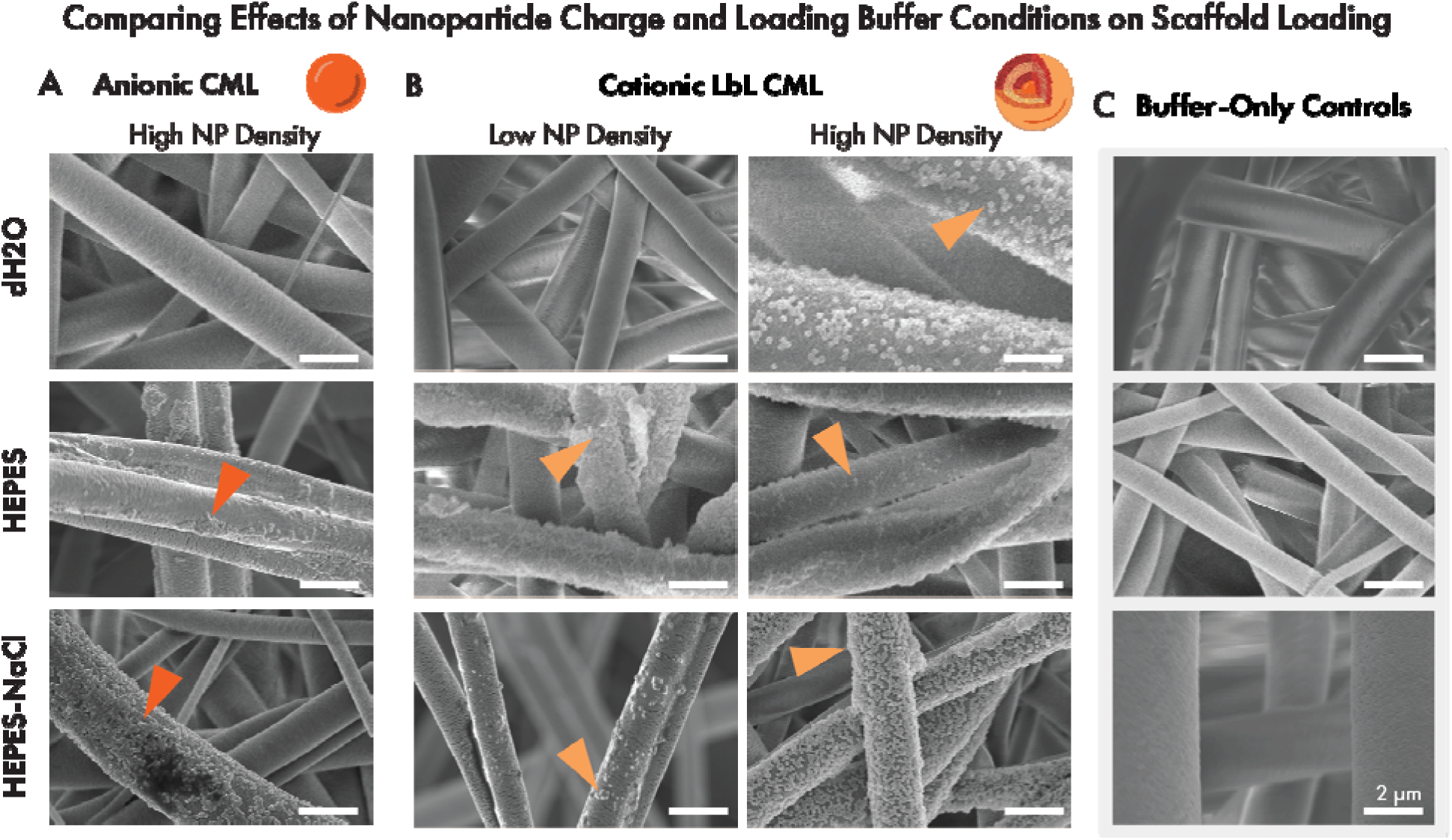
Solution conditions tune nanoparticle loading onto scaffolds. Scanning electron microscopy images of CML or LbL CML particle coatings at different densities (low=0.5 mg/mL and high=2 mg/mL) and in different loading buffers on electrospun scaffolds: (A) CML, (B) LbL CML, and (C) no particles. Arrows indicate discrete particles and/or particle plaques. Scale bar = 2 µm and is applied to all images.

Together, these results introduce a strategy for non-covalent, post-fabrication functionalization of piezoelectric scaffolds with nanotechnology. This is significant for tissue engineering applications, where electrospun fibers are commonly used as extracellular matrix (ECM) mimics.^26,61^ While strategies such as physical adsorption or encapsulation of proteins and small molecules,^62–65^ blend electrospinning of polymers with bioactive agents,^30,66–68^ coaxial electrospinning for core-shell architectures,^69–71^ and covalent immobilization of ligands or growth factors,^72–74^ have all been used to functionalize scaffolds, they often involve harsh processing, irreversible chemistry, or limited modularity.^75^ In contrast, our electrostatic adsorption method circumvents many of these issues by enabling post-fabrication modification under mild conditions, with tunable coating density and compatibility with both cationic and anionic nanoparticles. This makes it particularly suited for presenting fragile or multi-component nanocarriers and supports its utility in modular regenerative medicine platforms.

### 2.2 Extending nanomaterial coatings to liposomal nanotechnology

To adapt our nanoparticle coating platform for the delivery of biologically active signals, we selected liposomes as a modular and clinically relevant nanocarrier. Liposomes are biocompatible vesicles that can encapsulate small molecules within their bilayer or aqueous core, and their surfaces can be engineered to display proteins, lipids, or targeting ligands.^76–79^ Their surface chemistry, including overall charge, can be readily tuned by selecting different phospholipids and lipid components.^80^ By varying these parameters, liposomes can be engineered to support both electrostatic adsorption to the scaffold and programmable display of bioactive signals at the interface, making them a compelling platform for functionalizing scaffolds.

Building on our results with polymeric nanoparticles, we hypothesized that charged liposomes could similarly adsorb to the scaffold under optimized buffer conditions. We selected a cationic liposome formulation consisting of 1,2-distearoyl-sn-glycero-3-phosphocholine (DSPC), 1,2-dioleoyl-3-trimethylammonium-propane (DOTAP), and plant cholesterol, a 9:1:2 molar ratio, which was based on our previous work with liposomal biomaterials for drug delivery,^79^ and to promote adhesion to the negatively charged glycocalyx on mammalian cells.^81,82^ This design was intended to preserve scaffold-bound liposome accessibility and enhance cell–material interactions. We synthesized unilamellar liposomes with a mean diameter of 159.9 ± 0.3 nm, a low polydispersity index (0.05 ± 0.02), and a zeta potential of +50.75 ± 0.91 mV, indicating a stable, highly charged nanoparticle suitable for electrostatic self-assembly.^76,83^ (Figure 3A, Supplementary Figure 1).

**Figure 3.**
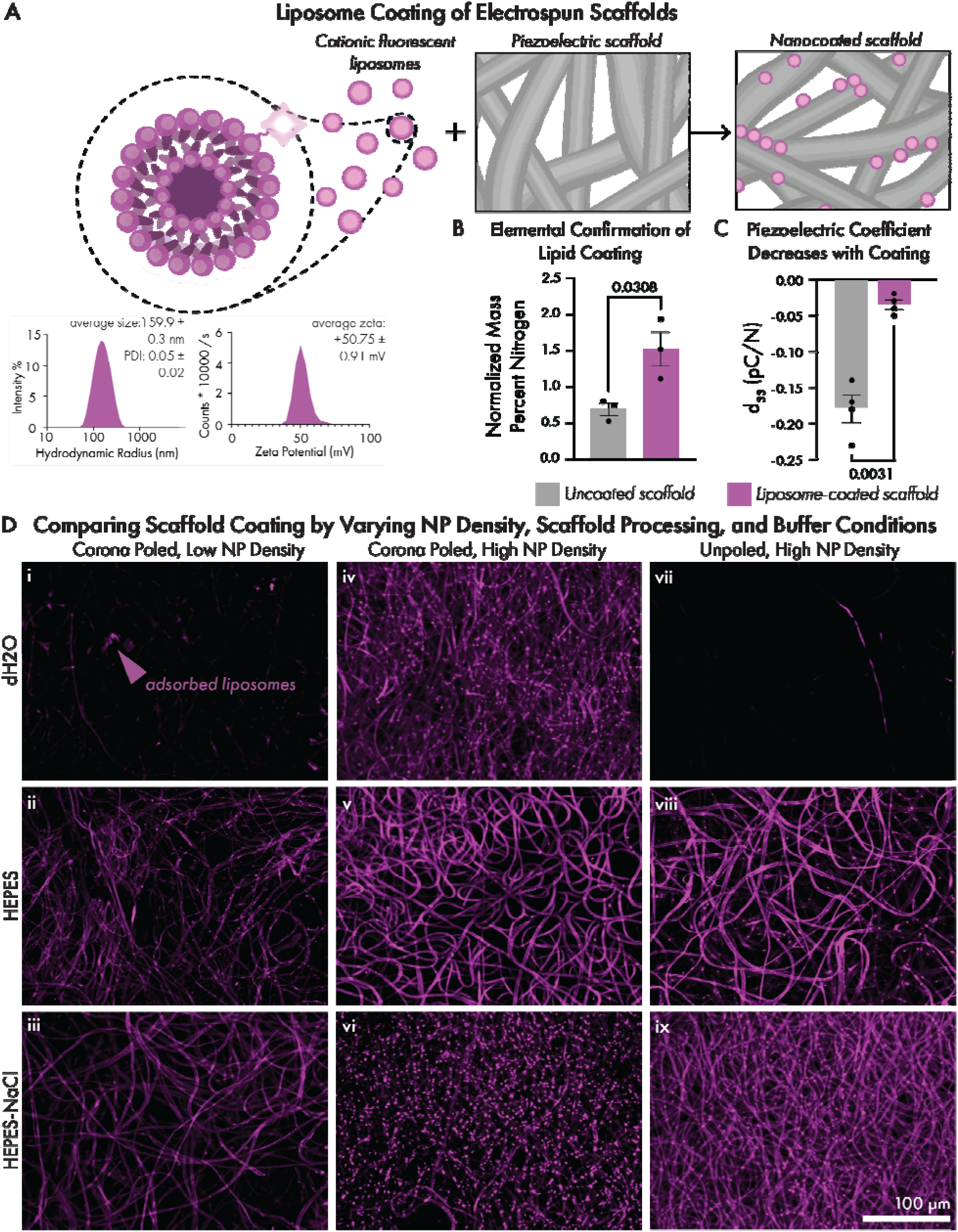
Electrostatic liposome adsorption onto PVDF-TrFE scaffolds: (A) Hydrodynamic radius and zeta potential of liposomes via dynamic light scattering: summary values are reported as mean ± SEM, with n=3 replicates. (B) Energy-dispersive X-ray spectroscopy measurements of nitrogen levels on coated vs. uncoated scaffolds, (C) piezoelectric coefficient, d_33_, on uncoated and coated scaffolds, and (D) fluorescence microscopy of Cy5 labeled liposomes on scaffolds coated in dH2O, HEPES, and HEPES-NaCl (top to bottom) at low NP density (0.5 mg/mL) or high NP density (2 mg/mL) on corona-poled scaffolds, or high NP density on a non-corona poled scaffold (left to right); scale bar =100 µm and is applied to all groups. Images are collected at the same laser power and are contrast-matched using Fiji-ImageJ. Pair-wise statistics are performed using student’s paired t-tests.

To initially validate liposome adsorption, we coated scaffolds using the optimal HEPES-NaCl buffer conditions we identified in the polymeric nanoparticle experiments. We used SEM combined with energy-dispersive X-ray spectroscopy (EDS) to detect levels of elemental nitrogen, which is present in the DSPC and DOTAP components of the liposome formulation, but is absent from the PVDF-TrFE scaffold itself. EDS confirmed that nitrogen levels were significantly higher in liposome-coated scaffolds compared to uncoated controls (1.522 ± 0.291 percent *vs.* 0.692 ± 0.107 percent, p value=0.03080), indicating successful adsorption (Figure 3B). We next asked whether nanoparticle coatings influenced the intrinsic piezoelectric properties of the scaffold. To assess this, we measured the piezoelectric coefficient (d_33_) of PVDF-TrFE mats before and after liposome coating. Coated scaffolds showed a significant (p=0.0031) reduction in piezoelectric response, with the magnitude of the d_33_ value decreasing from -0.18 ± 0.02 pC/N in uncoated scaffolds to -0.04 ± 0.01 pC/N after liposome adsorption (Figure 3C).

These findings suggest liposome adsorption may attenuate piezoelectric behaviors in the hybrid system.

To determine the tunability of liposome coatings, we asked whether the extent of adsorption could be modulated by solution conditions. This allowed us to test whether the design rules observed for polymeric nanoparticle coatings, specifically the role of pH and ionic strength, extend to lipid-based nanocarriers. We used the vacuum-assisted loaded technique to adsorb fluorescently labeled liposomes onto scaffolds using our three buffer conditions: deionized water, HEPES buffer (500 mM, pH=7.5), and HEPES (500 mM) further supplemented with NaCl (400 mM). Confocal microscopy revealed differences in coating density and uniformity that were consistent with our results with polymeric nanoparticles: the densest and most consistent coatings were observed in HEPES-NaCl, followed by HEPES alone, and scaffolds coated in deionized water displayed sparse and heterogeneous coating (Figure 3D). We also compared two concentrations of liposomes during loading: 0.5 mg/mL (Figure 3D i-iii) and 2 mg/mL of liposomes in loading buffer (Figure 3D iv-vi) and observed that coating density is proportional to liposome concentration. These findings confirm that pH stabilization, ionic strength, and liposome concentration can each influence adsorption outcomes, providing straightforward parameters for adjusting surface coverage.

We next investigated whether corona poling of the scaffold is required for liposome adsorption, or if optimized solution conditions and mechanical agitation during loading alone are sufficient. In deionized water, uniform liposome adsorption was only observed on corona-poled scaffolds (Figure 3 D iv), indicating that under unbuffered and low-ionic strength conditions, corona poling of the scaffold is required to create sufficient charge for adsorption. However, in HEPES-NaCl buffer, adsorption occurred efficiently on both poled and unpoled scaffolds (Figure 3 D vi, ix), suggesting that optimized solution conditions can compensate for the absence of corona poling during vacuum-assisted loading.

In contrast to the polymeric nanoparticle coatings, we observed a morphological difference in the liposomal coatings: corona poled scaffolds showed more punctate, clustered fluorescence (Figure 3D iv, vi), while unpoled scaffolds exhibited a more uniform coating (Figure 3D viii, ix). This difference was not observed in HEPES buffer without NaCl, where the coating morphology appeared similarly uniform between poled and unpoled scaffolds (Figure 3D v, viii). It is possible that in the presence of NaCl, the stronger or more spatially defined surface charges on poled scaffolds promote local liposome aggregation during adsorption, as inter-liposomal charge repulsion is reduced due to electrostatic screening. Overall, these findings suggest that transient charge domains generated by mechanical deformation are sufficient to support robust nanoparticle adsorption under optimal buffer conditions, potentially eliminating the need for time-intensive corona poling and allowing for a more efficient synthetic pipeline.

While liposomes adsorbed efficiently to PVDF-TrFE scaffolds under optimized conditions, several open questions remain. Corona poling was still necessary in low-ionic strength buffers and altered liposome coating morphology in the presence of salt, suggesting that scaffold polarization state may influence charge distribution and nanoparticle organization. The underlying mechanism remains unclear and could involve particle aggregation or localized differences in surface potential. We also observed a reduction in the scaffold’s piezoelectric coefficient following liposome coating. Although this is consistent with surface-level dielectric effects reported in other soft materials,^84,85^ it is not yet known how this change affects scaffold performance in biological contexts. Cells may engage with the scaffold by adhering, deforming fibers, or internalizing surface-bound materials, all of which could dynamically alter charge distribution or local piezoelectric behavior. Future studies will be needed to assess how surface coatings influence bioelectrical signaling during cell–material interactions.

### 2.3 Biocompatibility and immune cell engagement with liposome-coated scaffolds

To evaluate the biocompatibility of the liposome-coated scaffold system, we conducted *in vitro* studies with both HEK293 and Jurkat T cells. HEK293 cells were used as a standard model for assessing general cytocompatibility, while Jurkat T cells were used to explore the potential of this platform for *ex vivo* immune cell culture and applications within *in situ* cell therapy manufacturing. HEK293 cells were cultured on uncoated or cationic liposome-coated PVDF-TrFE scaffolds for eight days in low-adherence culture plates. Cell viability was quantified using the CCK8 assay, which measures metabolic activity via WST-8 reduction. HEK cells remained viable throughout the culture period, with no significant difference in metabolic activity between coated and uncoated scaffolds (p = 0.2300; Figure 4A). Jurkat T cells also remained viable when cultured on liposome-coated scaffolds, as confirmed by Calcein AM staining after three days in culture (Figure 4B).

**Figure 4:**
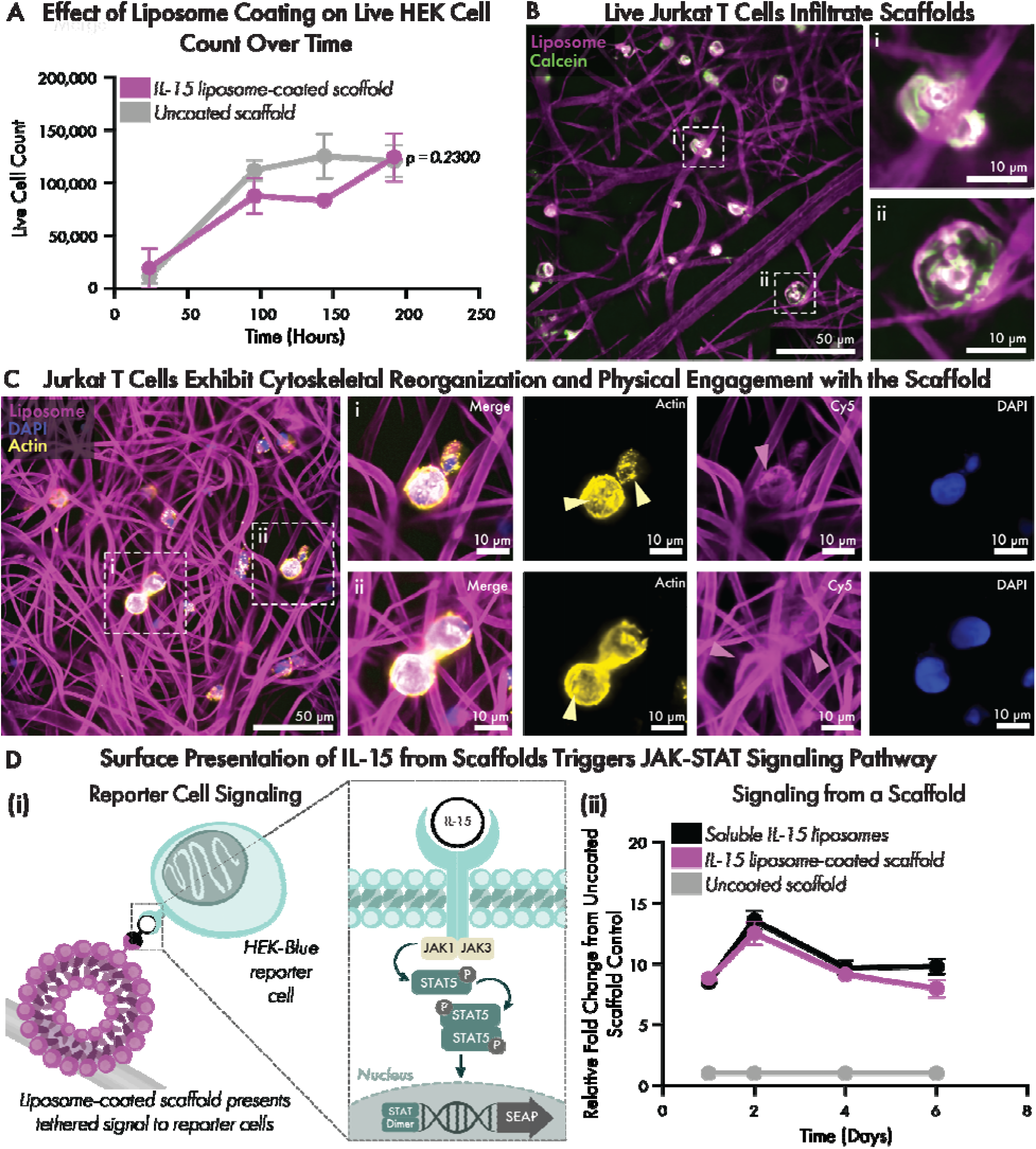
Liposome coated scaffolds are suitable substrates for cells and enable cytokine presentation: (A) Live HEK cell counts were measured from multiple time points on scaffolds with and without cationic liposome coating. Data is presented as mean ± SEM from 3 independent wells at each time point. Statistical significance was assessed by two-way ANOVA with Sidak’s multiple comparisons test, comparing coated to uncoated scaffold at each time point. P values reflect differences between groups at individual time points, and all error bars are present on graph although some are too small to see. (B) Scaffolds are coated with fluorescent Cy5-labeled liposomes and are imaged with confocal microscopy. Jurkat T cells are stained with Calcein, cultured on Cy5 liposome-coated scaffolds, and imaged at 60X resolution 3 days after plating. (C) Jurkat T cells are fixed on day 4 and stained intracellularly with ActinProbe555 and DAPI. Arrows indicate punctate actin (yellow) or liposome uptake (magenta). Scale bars are 50 µm or 10 µm. (D) Activating the JAK-STAT Pathway with IL-15 Liposomes on Scaffolds: (i) scaffolds are coated with IL-15-tethered liposomes in optimized HEPES-NaCl buffer conditions and SEAP is produced in response to IL-15 recognition. (ii) IL-15 reporter cells are plated on scaffolds, and absorbance is read daily, correlating to the downstream SEAP production from activation of the JAK-STAT pathway with n=3 samples per group. Data is presented as relative fold change from uncoated scaffold control, with mean ± SD. Significance is conducted with an Ordinary Two-Way Anova with multiple comparisons using Šídák’s multiple comparisons test.

To evaluate cell–material engagement, we examined cytoskeletal organization and assessed whether components of the liposome coating were transferred to the cell membrane. Jurkat T cells seeded onto Cy5-labeled liposome-coated scaffolds were fixed and stained for F-actin and DAPI on day four of cell culture. Confocal microscopy revealed that cells were adhered to scaffold fibers and exhibited a cortical actin distribution, with occasional actin-rich puncta at points of fiber contact (Figure 3C). In addition, Cy5-labeled lipids from the scaffold coating were detected within the plasma membranes of Jurkat T cells (Figure 4C, panels i,ii). This membrane-associated signal suggests that cells actively engage with and take up components of the scaffold’s liposome coating.

Unmodified PVDF-TrFE scaffolds have been used for tissue engineering,^30,86–88^ and our findings indicate that adding a cationic liposome coating preserves this biocompatibility while introducing a means for tunable nanomaterial presentation. Jurkat T cell behavior on these scaffolds was consistent with typical behaviors of immune cells interacting with three-dimensional substrates.^89^ Moreover, the incorporation of Cy5-labeled lipids into the cell membrane indicates active transfer of nanomaterials from the scaffold to cells, potentially through lipid exchange or fusion.^90,91^

### 2.4 Surface presentation of IL-15 cytokine from scaffolds supports functional JAK-STAT signaling

Having established that liposome-coated scaffolds support cell viability, we next investigated whether the scaffold interface could present functional signaling proteins in a localized, surface-bound format. In contrast to soluble cytokine delivery, which is often limited by rapid diffusion, degradation, and off-target effects, surface-tethered presentation enables more stable and spatially confined signal delivery.^92^ We have previously used histidine–nickel affinity interactions to tether recombinant cytokines to liposome surfaces in hydrogel systems,^79^ and we hypothesized that this supramolecular approach could be extended to liposomes adsorbed onto PVDF-TrFE scaffolds. To test this, we focused on interleukin-15 (IL-15), a cytokine of interest for *ex vivo* immune cell culture^93^ due to its role in promoting the survival and expansion of memory CD8+ T cells and NK cells.^10,94,95^

To tether IL-15 to the scaffold, we incorporated 0.3 mol%^79^ of the nickel-chelating lipid DGS-NTA(Ni) into the liposome bilayer and adsorbed the functionalized liposomes onto PVDF-TrFE scaffolds using the optimized vacuum-assisted loading protocol. His-tagged recombinant IL-15 was bound to the liposomes via incubation prior to coating liposomes onto scaffolds. To assess IL-15 bioactivity, we used HEK-Blue CD122/132 reporter cells, which secrete alkaline phosphatase (SEAP) in response to IL-15 receptor mediated phosphorylation of STAT5 (Figure 4Di). Reporter cells were seeded directly onto scaffolds in a non-TC-treated plate to encourage cell/scaffold integration, and SEAP levels were independently measured over the course of six days. We tested (i) IL-15 liposome-functionalized scaffolds, (ii) uncoated scaffolds with soluble IL-15, and (iii) uncoated scaffolds. STAT-5 phosphorylation kinetics increased similarly between cells cultured on scaffolds with soluble IL-15 or on scaffolds presenting liposome-bound IL-15 for days 1, 2, and 4 (p=0.9814, p=0.2453, p=8108, respectively) and only a moderate decrease in intensity was noted on day 6 (81.56% relative to soluble, p=0.0167) (Figure 4Dii). This reduction likely reflects minor IL-15 loss during coating, as the soluble control corresponded to 100% theoretical scaffold loading (0.477 µM IL-15).

These results confirm that scaffold-tethered IL-15 remains bioactive and capable of engaging cell surface receptors. More broadly, they demonstrate that liposome-coated scaffolds provide a modular platform for spatially confined cytokine presentation. By displaying IL-15 at the scaffold surface through supramolecular tethering, we preserved cytokine function and achieved localized receptor-mediated signaling without requiring covalent modification of either the cytokine or the scaffold. This flexible strategy enables facile incorporation of His-tagged proteins into electrospun materials and may prove particularly valuable for *ex vivo* immune cell expansion and reprogramming.

## Conclusion

This work establishes a modular strategy for post-fabrication functionalization of piezoelectric electrospun scaffolds using electrostatically adsorbed nanocarriers. By combining vacuum-assisted loading with buffer-optimized solution conditions, we enabled stable adsorption of charged nanoparticles, including both polymeric and lipid nanoparticles, onto PVDF-TrFE scaffolds. We applied this modular platform to a biomedically relevant context by generating scaffolds coated with liposomes decorated with immunomodulatory cytokines. Nanocoated scaffolds supported cell viability, enabled lipid exchange between immune cells and scaffold-bound liposomes, and activated cellular signaling pathways. Together, these findings demonstrate that the platform supports localized cytokine presentation and highlight its potential as an immune-instructive biomaterial in applications such as *ex vivo* 3D cell culture.

While this platform shows strong potential for modular signal presentation, several limitations remain that point to opportunities for future research. The ability to tether IL-15 to the scaffold surface and trigger localized signaling suggests this system could support immune-instructive cues in *ex vivo* or therapeutic settings. However, we evaluated cytokine activity using a HEK-based reporter line, and it remains to be determined whether the platform can drive similar or enhanced responses in primary immune cells such as T or NK cells. Finally, while our coating strategy preserves scaffold structure and supports cell viability, it is not yet clear whether the addition of a liposome layer affects the functional piezoelectric properties of PVDF-TrFE. Future studies using mechanosensitive readouts in relevant cell types, such as neurons,^86^ could help determine whether bioelectric signaling is maintained alongside biochemical signaling in this system.

Looking ahead, this electrostatic coating strategy could be extended to a broad range of surface-charged nanocarriers, including extracellular vesicles, polymeric micelles, hybrid nanoparticles, and solid lipid nanoparticles (LNPs). Incorporating LNPs would enable scaffold-tethered gene delivery, combining nucleic acid cargo with the structural and biochemical advantages of electrospun materials. The use of liposomes as a modular display vehicle also opens opportunities for multivalent or combinatorial presentation of proteins, such as co-delivered cytokines, chemokines, and growth factors. More broadly, the ability to tune surface coverage through buffer composition and nanoparticle concentration offers a simple and scalable method for customizing scaffold interfaces. Together, these features position this approach as a flexible foundation for engineering multifunctional, cell-responsive scaffolds with potential applications in regenerative medicine, immune engineering, and *ex vivo* cell manufacturing.

## Materials and methods

### PVDF-TrFE Scaffold Fabrication

Scaffolds are electrospun by modifying previously reported methods.^47,86^ Briefly, 20% (w/v) solutions of Polyvinylidene fluoride tri-fluoroethylene (PVDF-TrFE, 70/30) (400kDa, poly-dispersity index (PDI) of 2.1, Solvay Solexis Inc., Thorofare, NJ) in 2-Butanone/methyl ethyl ketone (Sigma-Aldrich # 360473) are filled into a 10mL syringe and extruded though an 18-gauge needle, with a 3ml/hour flow rate. A voltage of 22 kV is connected to the needle tip and randomly oriented fibers are collected on a grounded metal collector plate placed 22-24 cm apart from the needle tip. The fibers are spun at room temperature (∼20°C) with 20-30% humidity followed by aerating the electrospun fibrous mat for at least 48 hours. The scaffolds are annealed at 135°C for 96 hours and quenched as described in previous methods.^30^

### Corona poling

The scaffolds are corona poled using a custom setup described previously.^47^ Briefly, the scaffolds are heated to a temperature of ∼47°C for 3 hours on a grounded heating plate. The positive charge is deposited on the scaffold surface using needle tips connected to a voltage of 14 KV. A metal grid with 2 KV voltage is set 2 cm above the scaffolds to distribute the positive charges.

Scaffold fiber diameter measurements: A total of 40 fibers from different spots of 5 distinct SEM images were quantified using Fiji-Image J. Measurements are reported as mean ± SD.

### Layer-by-Layer Nanoparticle Synthesis

100 nm Carboxyl-modified latex nanoparticles (CML) (ThermoFisher Scientific # F8801) are coated in three layers of poly-allylamine hydrochloride (PAH) (Sigma-Aldrich # 283215), poly(acrylic acid, sodium salt) solution (PAA) (Sigma-Aldrich # 416029) and PAH. PAH is prepared at 4 mg/mL and 6 mg/mL in MilliQ water. PAA is prepared at 4 mg/mL in MilliQ water. CML are prepared at 1 mg/mL and are combined with equal volume of PAH. The solution is pipetted in a bath sonicator three times. The particles are spun at 25,830 x g for 45 minutes at 4°C. The supernatant is removed, and 0.8 times the original particle volume minus the particle volume remaining of MilliQ water is added to the solution to resuspend the particles. Equal volume of 4 mg/mL PAA is added to the particle solution by pipetting inside a bath sonicator. The particles are spun at 25,830 x g for 45 minutes at 4°C. The supernatant is again removed, and 0.8 times the original particle volume minus the particle volume remaining of dH2O is added to the solution to resuspend the particles. Equal volume of 6 mg/mL PAA is added to the particle solution by pipetting inside a bath sonicator. The particles are spun at 25,830 x g for 45 minutes at 4°C. The supernatant is removed, and the particles are resuspended in loading buffer.

### Liposome Synthesis

Liposomes are prepared using the thin-film rehydration technique: 1,2-distearoyl-sn-glycero-3-phosphocholine (18:0 PC (DSPC)) (Avanti Polar Lipids # 850365), 1,2-dioleoyl-3-trimethylammonium-propane (chloride salt) (18:1 TAP (DOTAP)), (Avanti Polar Lipids # 890890), cholesterol (plant) (Avanti Polar Lipids # 700100), and 1,2-dioleoyl-sn-glycero-3-[(N-(5-amino-1-carboxypentyl)iminodiacetic acid)succinyl] (nickel salt) (18:1 DGS-NTA(Ni)) (Avanti Polar Lipids # 790404) are dissolved at a 9:1:2:0.379 molar ratio in chloroform. Solvent is removed by evaporation under vacuum using a rotovap with bath at 65 °C. The thin film is rehydrated at the desired concentration in loading buffer by in a bath sonicator at 60°C for 30 mins. For fluorescent liposomes only, 1,2-distearoyl-sn-glycero-3-phosphoethanolamine-N-(Cyanine 5) fluorescent lipid (18:0 Cy5 PE) (Avanti Polar Lipids # 810345) is added at 0.01 mol% during sonication. Liposomes are extruded at 60°C (Avanti Research # 610000) using 10 mm filter supports (Avanti Polar Lipids # 610014) through PC 1.0 µm membrane (Avanti Research # 610010), PC 0.4 µm membrane (Avanti Research # 610007), PC 0.2 µm membrane (Avanti Research # 610006) and PC 0.1 µm membrane (Avanti Research # 610005).

### Nanoparticle Characterization

Nanoparticle hydrodynamic radius and polydispersity index were measured using dynamic light scattering (Malvern Zetasizer Advanced Pro Red, λ=633 nm, dispersant RI=1.33. Zeta potential measurements were also acquired with the Malvern Zetasizer, using laser Doppler electrophoresis. Nanoparticles were diluted 1:1000 in Milli-Q water, with no added salts, in polystyrene Greiner Bio-One Semi-micro/Macro cuvettes (Fisher Scientific # 07-000-571) or Folded Capillary Zeta Cell (Malvern Paranalytical # DTS1070) for characterization.

### Scaffold Loading

Loading buffers were formulated in MilliQ water: HEPES-NaCl (500 mM HEPES 400 mM NaCl) and 400 mM NaCl; HEPES Sodium Salt (White Powder), Fisher BioReagents (Fisher Scientific # BP410-500), Sodium Chloride (Fisher Scientific # BP358-1). For all experiments, the scaffold had a size of 6 mm diameter x 0.35 mm thickness, which were cut from the electrospun mat using a biopsy punch (Integra # 33-36). For scaffold loading by incubation, the scaffold disks were placed in a round-bottom 96-well plate. 200 uL of nanoparticle solution at the desired concentration was added and scaffolds were incubated for 1 hour, after which they were washed three times with MilliQ water. For vacuum-assisted loading, 6 mm scaffold disks were transferred into separate 5 ml polypropylene round-bottom tubes (Corning # 352063). 265 µL of the nanoparticle mixture at the desired concentration in loading buffer was added to each tube and capped. A 21-gauge needle (Sigma-Aldrich # Z192503) attached to a 5 mL syringe was inserted into the cap of the tube and was used to apply vacuum by pulling back on the plunger and then, agitation was applied for 30 seconds. This process was performed twice. The tubes were then incubated for one hour at 4°C, after which the mixture was aspirated, and the scaffolds were rinsed three times with DI water. All scaffolds used in these experiments were from the same batch of electrospun scaffold mat.

### Scaffold Imaging

The CML and LbL CML nanoparticle-loaded scaffolds were air dried overnight prior to SEM sample preparation. The samples were 8-10 nm sputter-coated with gold-palladium (Cressington sputter coater- 108 auto) and imaged using Zeiss Sigma VP SEM with an accelerating voltage of 5 KV. At least n=2 images were captured on two different areas from each scaffold with approximately 1500X and 7000X magnification.

Fluorescent liposome-coated scaffolds were imaged using the Nikon Ti2 inverted microscope with AXR resonant spectral scanning confocal unit with laser emission range 662nm-737nm. Scaffolds were prepared on a microscope slide and imaged through a coverslip. Z-stacks of 50 µm were acquired to create maximum intensity projections. Images were denoised and processed with FIJI-ImageJ software.

### Elemental Analysis

Energy dispersive spectroscopy (EDS) was used for quantitative analysis of liposome-coating. Liposome-coated scaffolds were dried for 48 hours prior to analysis. Samples were coated with 10nm carbon and imaged using the Screening Cube II (SRAMM). EDS measurements were collected with 20,000 counts at approximately 8100X magnification. n=2 samples were prepared for each group, and an unpaired student’s t-test was performed.

### Piezoelectric Coefficient (d33) Measurements

Annealed scaffolds were cut into 12 mm diameter x 0.35 mm thickness disks and the piezoelectric coefficient was measured with a piezometer (Piezotest-PM300). The scaffolds were then coated with liposomes using our vacuum-assisted loading technique and airdried overnight. Piezoelectric coefficient measurements were repeated 24h later for the liposome coated scaffolds and uncoated experimental control scaffolds.

### Cell Culture

HEK Blue™ CD122/CD132 (Invivogen # hkb-il2bg) cells were maintained in Dulbecco’s Modified Eagle’s Medium (DMEM) (Corning # 10-014-CM) supplemented with 10% heat-inactivated fetal bovine serum (Fisher Scientific # A5256701), 100U/mL penicillin, 100µg/mL streptomycin (Thermo Fisher Scientific # 15140122) 2 mM L-Glutamine (Millipore Sigma # G7513), 100 µg/mL Normocin™ (Invivogen # ant-nr), supplemented with 1X HEK-Blue™ Selection (Invivogen # hb-sel) and 1 µg/mL Puromycin (Invivogen # ant-pr) to select for reporter cells. Cells are tested for mycoplasma and are maintained at 37°C and 5% CO2.

### Cell Viability

Cationic liposomes were formulated in sterile HEPES-NaCl (500 mM HEPES 400 mM NaCl) buffer. Scaffolds were loaded with liposomes or sterile buffer via vacuum-assisted loading for each time point. HEK Blue™ CD122/CD132 (Invivogen # hkb-il2bg) cells were plated on scaffolds, and this procedure was repeated every other day, in addition to replacing the cell media in each scaffold well, to enable a comparable readout at the end timepoint. Eight days after plating the first timepoint, Apexbio Technology LLC Cell Counting Kit-8 (CCK-8) reagent (Fisher Scientific # 50-190-5565) was added to the scaffold-cell solution achieve a final concentration of 10% CCK-8 reagent in the cell culture well. A standard curve was generated to interpolate cell count values using simple linear regression. The plate was agitated and incubated for three hours at 37°C, at which point the plate was agitated again in the Synergy Neo2 Plate reader, scaffolds were removed, and endpoint absorbance was read at 450 nm.

### Cell Signaling Assay

Scaffolds are punched using 6mm biopsy punches (Integra Life Sciences # 33-36) and UV-treated for 15 mins on each side. Liposomes are incubated with recombinant Histidine-tagged human IL-15 (Sino Biological # 10360-H07E) overnight at 4°C to achieve a concentration of 180 ng/mL IL-15 on liposomes at 2 mg/mL, the loading concentration, in HEPES-NaCl. Scaffolds are coated with IL-15 liposomes as previously described with sterile vacuum-assisted loading techniques and are placed in flat-bottom propylene non-TC treated 96-well plates (Fisher Scientific cat# 07-200-696).

Scaffolds are incubated with complete media overnight at 4°C. Media is removed, and 50,000 HEK-Blue CD122/CD132 (Invivogen # hkb-il2bg) cells at passage 8 are added to the liposome-coated scaffolds, uncoated scaffolds, or soluble condition (containing 0.477 µM of IL-15, the amount of 100% loading efficiency of IL-15 on scaffolds) in 100 uL complete media without Normocin or selective antibiotics. Every 24 hours, the media is collected to be stored at 4°C. One week after plating, QUANTI-Blue™ solution (Invivogen # rep-qbs) is prepared and incubated at a 10:1 ratio with collected media for 30 minutes at 37°C. SEAP levels are obtained with endpoint absorbance readings using the Synergy Neo2 plate reader at 620 nm.

## Acknowledgements

This work was funded by Columbia University’s Provost’s Cluster Hire Pilot Award in the Area of STEM Research Support. SSC and TA are funded by the National Science Foundation Science and Technology Center – Center for Engineering Mechanobiology (CEMB), CMMI – 1548571. These studies used the Confocal and Specialized Microscopy Shared Resource of the Herbert Irving Comprehensive Cancer Center at Columbia University, funded in part through the NIH/NCI Cancer Center Support Grant P30CA013696. Some of this work was performed at the Simons Electron Microscopy Center at the New York Structural Biology Center, with major support from the Simons Foundation (SF349247). The authors acknowledge the use of facilities, including Zeiss Sigma VP SEM, supported by NSF through the Columbia University, Columbia Nano Initiative, and the Materials Research Science and Engineering Center DMR-2011738.

The authors would like to acknowledge Yasaman Aghli and Anna Kylat for contribution on fabricating electrospun scaffold mats and providing materials for in vitro assays.

## Author Contributions

Sarah Payne Bortel (**SB**), Sumayia Saif Jaima Chowdhury (**SSC**), Jeremy Cheng (**JC**), Daniella Uvaldo (**DU**), Mackenzie Wright (**MW**), Treena Livingston Arinzeh (**TA**), Santiago Correa (**SC**). **SB**: Conceptualization, data curation, formal analysis, investigation, methodology, supervision, visualization, validation, writing – original draft. **SSC**: conceptualization, data curation, investigation, methodology, validation. **JC**: data curation, investigation, validation, methodology. **DU**: investigation, validation, methodology. **MSW**: investigation, validation, methodology. **TA**: conceptualization, funding acquisition, project administration, resources, supervision. **SC**: conceptualization, funding acquisition, project administration, resources, supervision, writing – review & editing.

