## Supplementary Information for "Modular Noncovalent Functionalization of Electrospun Piezoelectric Scaffolds with Bioactive Nanocarriers"

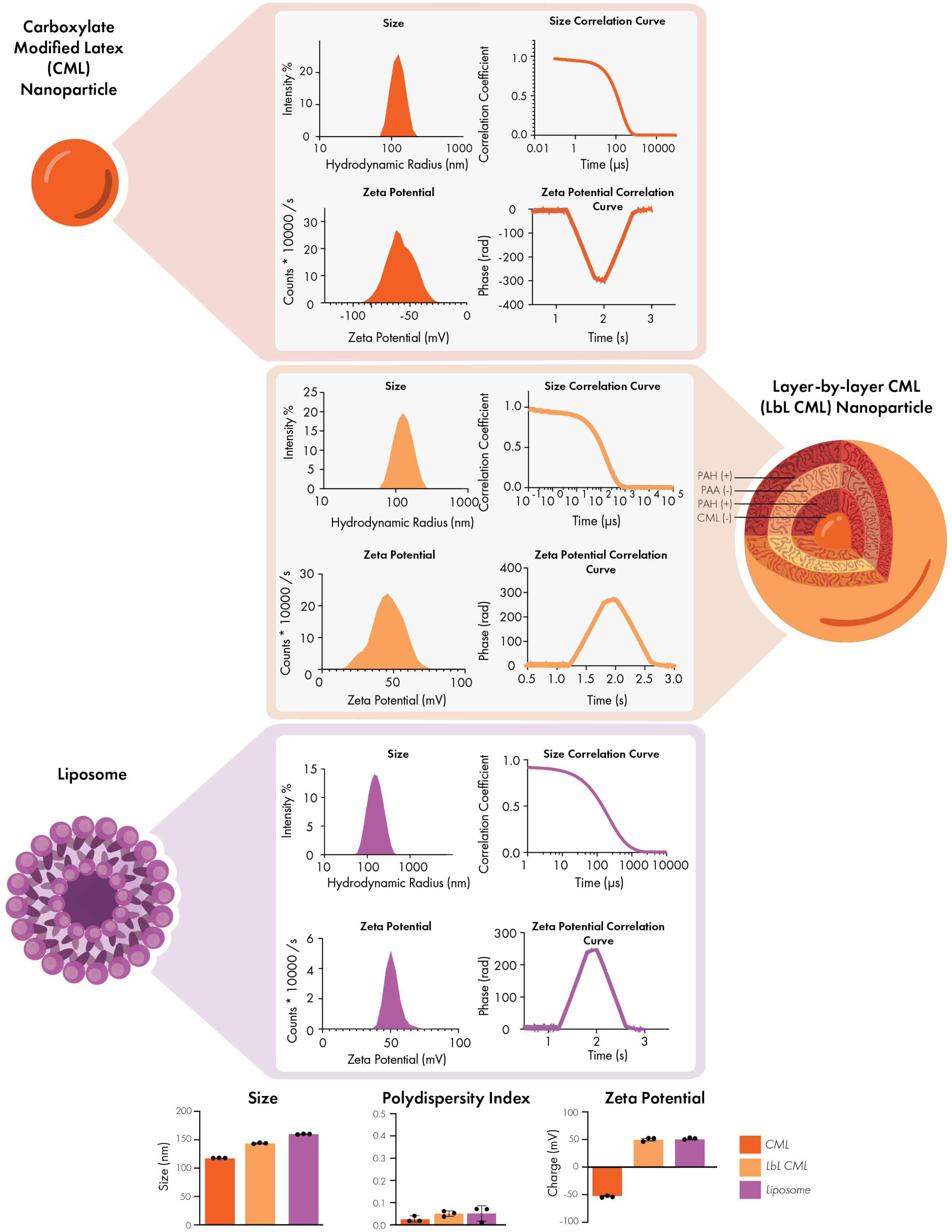


**Supplementary Figure 1: Nanoparticle summary data.** Dynamic light scattering is used to characterize nanoparticle size, polydispersity index, and zeta potential. Carboxylate modified latex (CML) particles (top), layer-by-layer CML (LbL CML) (middle), and liposomes (bottom) are shown. Results are reported as the mean of n=3 independent syntheses. Individual mean data points are plotted blow, and results are represented as mean ± SEM.

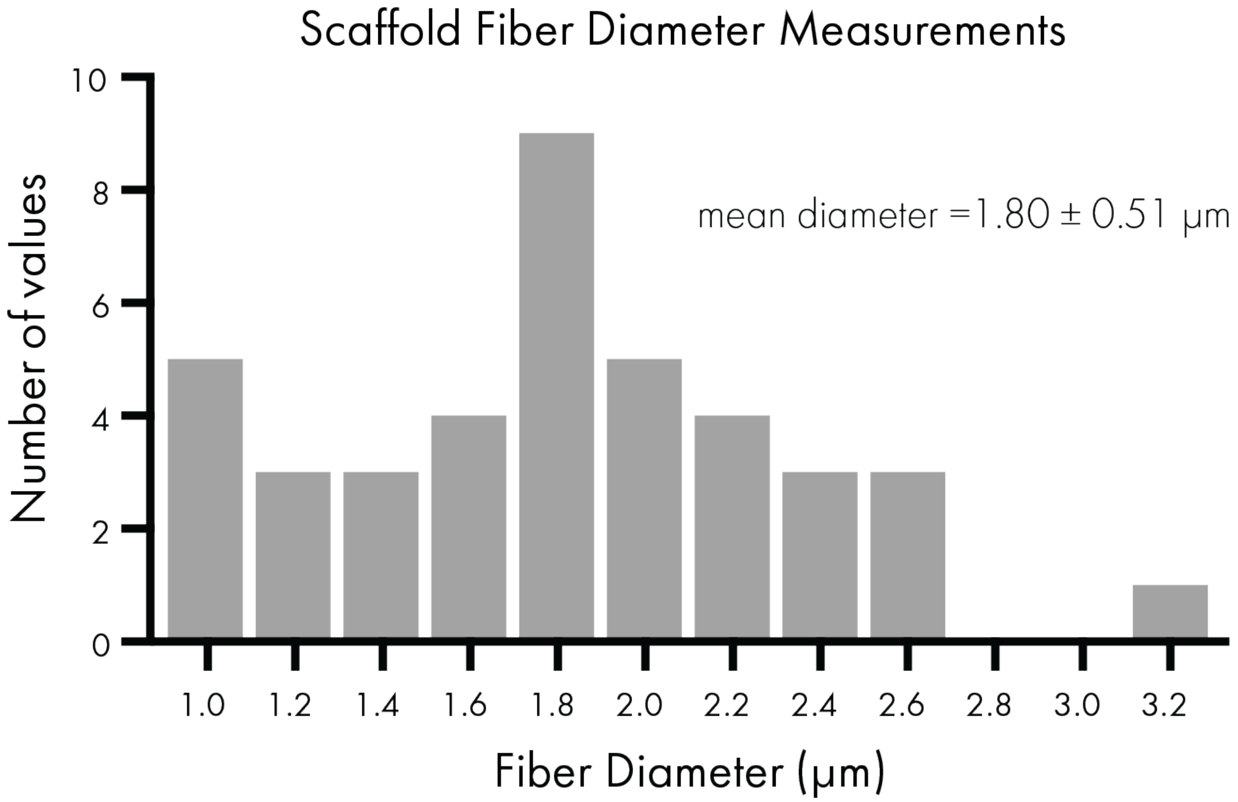


**Supplementary Figure 2: Scaffold fiber size distribution.** 40 fiber measurements are collected using Fiji ImageJ from 5 independent scanning electron microscopy images, and a histogram of the frequency distribution is shown. Summary result of fiber diameter is reported as mean ± SD.
